# Retinoic acid and proteotoxic stress induce AML cell death overcoming stromal cell protection

**DOI:** 10.1101/2023.02.20.529204

**Authors:** Francesca Liccardo, Martyna Śniegocka, Claudia Tito, Alessia Iaiza, Tiziana Ottone, Mariadomenica Divona, Serena Travaglini, Maurizio Mattei, Rossella Cicconi, Selenia Miglietta, Giuseppe Familiari, Stefania Annarita Nottola, Vincenzo Petrozza, Luca Tamagnone, Maria Teresa Voso, Silvia Masciarelli, Francesco Fazi

**Author notes:** corresponding authors: Francesco Fazi, PhD,; Silvia Masciarelli, PhD.

## Abstract

Acute myeloid leukemia (AML) patients bearing the ITD mutation in the tyrosine kinase receptor FLT3 (FLT3-ITD) present a poor prognosis and a high risk of relapse. FLT3-ITD is retained in the endoplasmic reticulum (ER) and generates intrinsic proteotoxic stress. We devised a strategy based on proteotoxic stress, generated by the combination of low doses of the differentiating agent retinoic acid (R), the proteasome inhibitor bortezomib (B), and the oxidative stress inducer arsenic trioxide (A). It exerts strong cytotoxic activity on FLT3-ITD^+^ AML cell lines and primary blasts isolated from patients, due to ER homeostasis imbalance and generation of oxidative stress. AML cells become completely resistant to the combination RBA when treated in co-culture with bone marrow stromal cells (BMSC). Nonetheless, we could overcome such protective effects by using high doses of ascorbic acid (Vitamin C) as an adjuvant. Importantly, the combination RBA plus ascorbic acid significantly prolongs the life span of a murine model of human FLT3-ITD^+^ AML without toxic effects. Furthermore, we show for the first time that the cross-talk between AML cells and BMSC upon treatment involves disruption of the actin cytoskeleton and the actin cap, increased thickness of the nuclei, and relocalization of the transcriptional co-regulator YAP in the cytosol of the BMSC. Our findings strengthen our previous work indicating induction of proteotoxic stress as a possible strategy in FLT3-ITD^+^ AML therapy and open to the possibility of identifying new therapeutic targets in the crosstalk between AML cells and BMSC, involving mechanotransduction and YAP signaling

## Introduction

Acute myeloid leukemia (AML) is a heterogeneous disease caused by a blockage of hematopoietic myeloid precursors differentiation, resulting in the accumulation of immature blasts. The mutational landscape of AML comprises mutations in signaling pathways, transcription factors, epigenetic modifiers, and splicing factors^1^. FMS-like tyrosine kinase 3 (FLT3) internal tandem duplication (FLT3-ITD) and tyrosine kinase domain (FTL3-TKD) mutations cause constitutive activation of FLT3 and its downstream signaling pathways^2–5^. FLT3-ITD mutations are associated with a poor prognosis, primarily due to an increased risk of relapse, with a median overall survival (OS) at five years of 30-35 %^6,7^. The clinical outcome of FLT3-ITD AML and the strong evidence of the leukemogenic role of FLT3, promoted the development of tyrosine kinase inhibitors (TKis)^8,9^. However, despite encouraging results achieved with targeted treatments, in association or not with chemotherapy, improvements in overall survival of FLT3-ITD^+^ AML are still modest, since most patients eventually relapse^10–13^. Acquisition of resistance to treatments involves the development of FLT3 secondary mutations^14^, the evolution of leukemic clones characterized by novel mutations (especially involving the RAS pathway^15^), and protection provided by the bone marrow (BM) niche^16,17^. These observations suggest that a possible strategy is to combine multiple approaches targeting different pro-survival pathways. Leukemic cells rely on adaptive responses to cope with intrinsic and extrinsic sources of proteotoxic stresses, that result in the disruption of protein homeostasis^18^. In this context, we set up an approach based on the induction of endoplasmic reticulum (ER) and oxidative stress that showed cytotoxic activity against FLT3-ITD^+^ AML, both in cell lines and primary blasts isolated from patients^19^. This is based on the combined use of low doses of the differentiating agent retinoic acid (RA), the ER stress inducer tunicamycin (Tm), and the oxidative stress inducer arsenic trioxide (ATO) and importantly did not show toxicity on normal hematopoietic progenitors treated *ex vivo*. To facilitate possible translational applications, here we substituted Tm with the proteasome inhibitor bortezomib (Btz) to induce ER stress and alteration of proteostasis. Indeed, RA and ATO are employed in clinical practice for acute promyelocytic leukemia^20^ and Btz for multiple myeloma and mantle cell lymphoma^21,22^. We demonstrate that the combination of RA, Btz and ATO (RBA) efficiently targets FLT3-ITD^+^ AML cells. Furthermore, since the BM niche plays a crucial role in protecting leukemic cells from the cytotoxic effects of different therapies^16,17^, we assessed the efficacy of the combination RBA on AML cells in a coculture system with BM stromal cells (BMSC). We found that BMSC render AML cells totally resistant to the combination RBA, but the addition of ascorbic acid (Vitamin C) is sufficient to reduce the protective effects. We also investigated the molecular basis involved in the cross-talk between AML cells and BMSC upon treatment demonstrating the involvement of actin cytoskeleton dynamics and of the transcriptional co-regulator YAP. Eventually, we tested the efficacy of the combination RBA plus ascorbic acid in a murine model of human FLT3-ITD^+^ AML, and found that it significantly prolongs the mice life span, reducing AML engraftment.

## Methods

An extended Methods section is available in the supplemental material.

### Cell culture

Human AML cell lines were cultured in RPMI 1640 medium supplemented with 1% penicillin/streptomycin and 10% FBS. Murine BMSC line MS-5 was cultured in MEMα with ribonucleosides and deoxyribonucleosides, with 1% penicillin/streptomycin, 20% FBS and βmercaptoethanol 100μM.

### Patient samples

AML BM samples were collected at diagnosis at the Department of Biomedicine and Prevention at the University of Rome Tor Vergata, after obtaining informed consent from all patients and approval of the study by the IRB of Policlinico Tor Vergata, Rome. After purification CD34+ LSC were cultured *in vitro* in STEM SPAN Leukemic cell expansion medium.

### Treatments

MOLM-13 cells were treated with 10nM RA, 2.25nM Btz and 500nM ATO, alone or in combination, as described. In the co-culture system, ascorbic acid was added at a 4.5mM concentration.

### Measurement of cell death and cell proliferation

We performed flow cytometry to assess cell death and cell cycle.

### Evaluation of ER and oxidative stress

We performed RT-qPCR, flow cytometry, confocal microscopy, and TEM analysis^23,24^ to evaluate the activation of the UPR and of the anti-oxidant response upon treatments.

### Analyses of the co-culture system

We performed flow cytometry, RT-qPCR, confocal microscopy, and WB analysis to investigate the responsiveness of the co-cultures to the treatments.

### *In vivo* mouse studies

Orthotopic human leukemia was induced by injecting MOLM-13 cells in NSG mice. The study was approved by the Italian Ministry of Health.

## Results

### The combination of low doses of Retinoic Acid, Bortezomib and Arsenic Trioxide induces cell death of FLT3-ITD^+^ AML cell lines and primary human leukemic stem cells

We screened a panel of human AML cell lines to assess their sensitivity to the combination of RA, Btz, and ATO (RBA). As we observed in our previous work^19^, the cell lines expressing FLT3-ITD resulted to be the most affected by the induction of proteotoxic stress (Supplemental Table 1). We chose doses of each drug that did not cause significant cell death or cell cycle alterations when used alone, but only in combination. The triple combination produced the highest cytotoxic effect on the FLT3-ITD^+^ MOLM-13 cells (Figures 1A, B, and C) as well as on MOLM-14 and MV-4-11 cell lines (FLT3-ITD^+^ as well, data not shown). We then tested the combination on FLT3-ITD^+^ primary AML leukemic stem cells (CD34^+^) and confirmed its highest efficacy, even though these resulted quite sensitive to the double combinations as well (Figure 1D).

**Figure 1.**
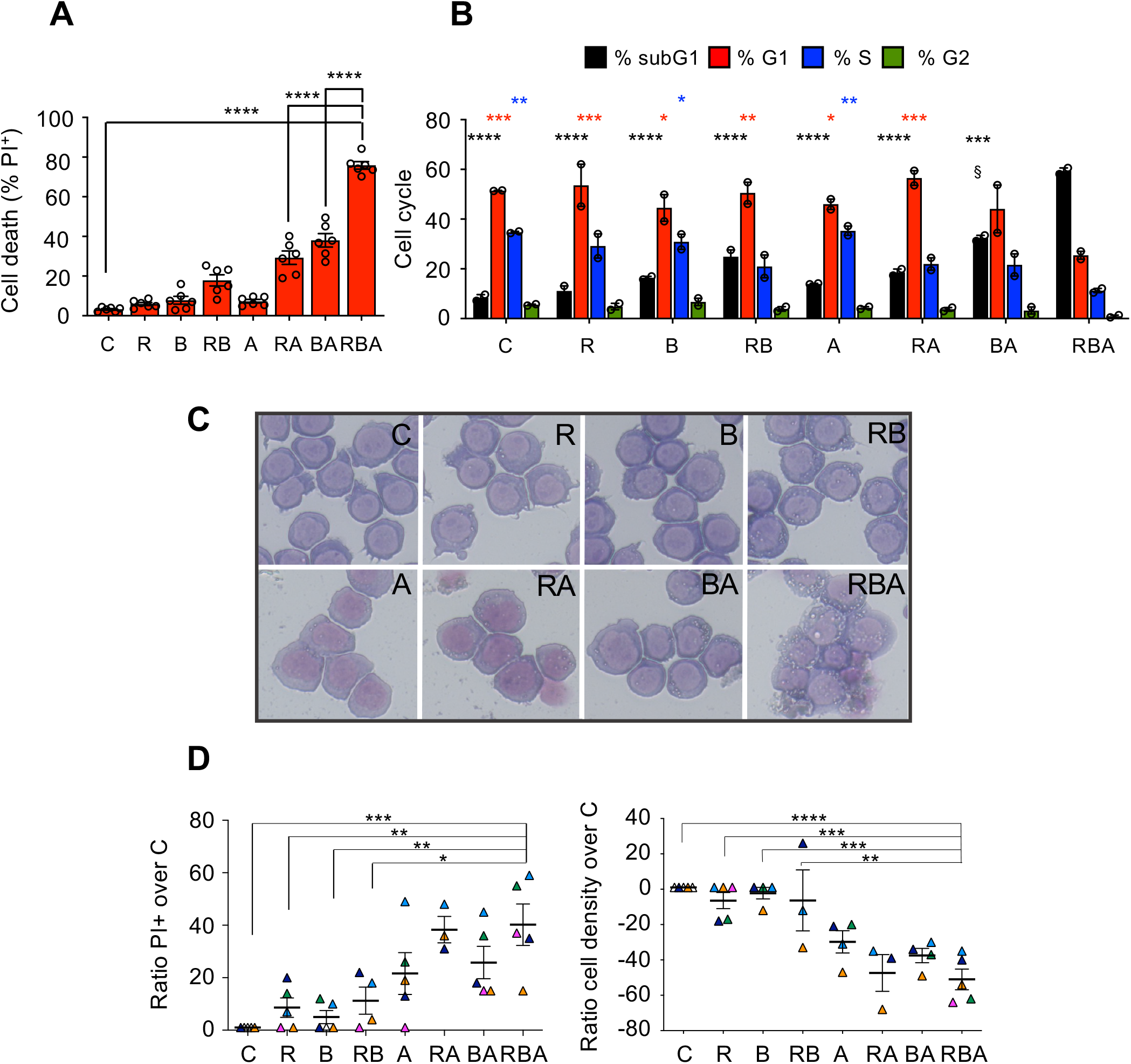
The FLT3-ITD^+^ MOLM-13 cells and primary leukemic stem cells are sensitive to the combination of RA, Btz and ATO. **A)** MOLM-13 AML cells were treated for 72 hours with 10nM RA, 2.25nM Btz, and 500nM ATO, alone or in combination. In all the figures retinoic acid is indicated as R, bortezomib as B, and ATO as A. Cell death was assessed by propidium iodide (PI) exclusion assay, analyzed by flow cytometry (one-way ANOVA). **B)** Cell cycle analysis of MOLM-13 cells 48 hours after treatments as in (A). The cells treated with BA show an increased percentage of subG1 phase relative to C cells but only those treated with RBA exhibit significant modulation of all the cell cycle phases. (two-way ANOVA vs RBA; § subG1 BA vs subG1 C: P <0.005). **C)** MOLM-13 cell morphology analyzed by Wright-Giemsa staining, 72 hours after treatments. **D)** FLT3-ITD^+^ CD34^+^ cells, isolated from patients at diagnosis, were treated *ex vivo* for 96 hours with 10nM RA, 3nM Btz, and 500nM ATO, alone or in combination. Cell death (left) and cell density (right) were measured by flow cytometry after staining with PI (one-way ANOVA).

### Treatment with the combination RBA alters ER homeostasis in MOLM-13 cells but the adaptive branch of the UPR is suppressed

We expected that the combination RBA would cause proteotoxic stress because the three drugs affect ER functions and protein folding. Although AML subtypes like MOLM-13 cells do not respond to the differentiation stimulus provided by RA in such a complete manner as acute promyelocytic leukemia cells do^25^, RA could still induce partial differentiation by increasing ER activity. Indeed, we observed activation of the secretory pathway upon RA administration, as indicated by diminished nucleus/cytosol ratio and cytosolic basophilia (Figure 1C), swollen ER tubules (Figure 2A), and augmented expression of the ER chaperones calnexin and BiP (Figure 2B). Since Btz inhibits the 26S subunit of proteasome, it blocks the final step of the ER-associated degradation (ERAD), the process through which misfolded proteins accumulated in the ER are eventually degraded^26^. Thus, by administrating Btz we expected to induce proteostasis unbalance, affecting ER protein folding capacity. Furthermore, ATO induces oxidative stress, which impairs protein folding^27^. Indeed, MOLM-13 cells treated with RBA present swollen ER tubules (Figure 2A) and increased expression of the ER chaperone calnexin (Figure 2B), indications of increased activity of the ER. ER stress is defined as the accumulation of misfolded/unfolded proteins in the ER lumen exceeding the folding capacity of the ER. Such circumstances trigger a series of signaling pathways, collectively known as the unfolded protein response (UPR), which regulate the expression of genes involved in protein folding and secretion to re-establish ER homeostasis. However, in case of prolonged or overwhelming stress, the UPR leads to apoptotic cell death^28^. Surprisingly, despite increased ER activity, MOLM-13 cells treated with RBA not only do not up-regulate the expression of the two main UPR players involved in homeostasis recovery, namely the ER chaperone BiP and the spliced form of the transcription factor XBP1, but this is down-regulated, further worsening ER proteostasis unbalance, whereas the pro-apoptotic UPR gene CHOP is not modulated (Figure 2C and Supplemental Figure 1). The same down-regulation of BiP and sXBP1 is observed upon administration of ATO, alone or in combination with RA or Btz, indicating induction of proteotoxic stress.

**Figure 2.**
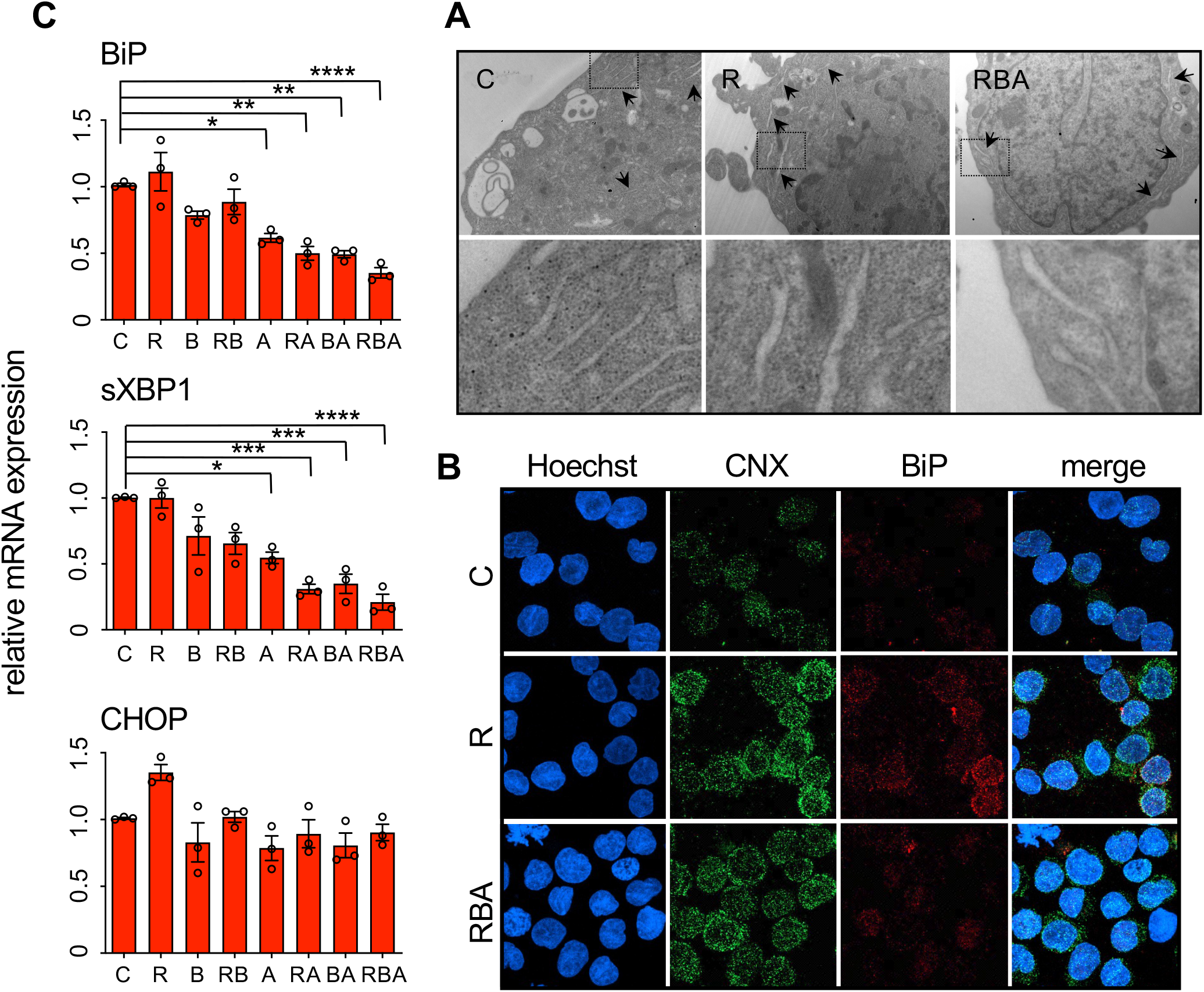
RBA combined treatment generates ER stress without consequent activation of the pro-survival UPR pathway. **A)** TEM analysis of MOLM-13 cells, 24 hours after treatment, shows swollen ER in the cells treated with R alone and with the combination RBA relative to C cells. Black arrows point to ER tubules. Dashed squares in the upper panels indicate the areas magnified in the lower panels. **B)** Confocal microscopy images of MOLM-13 cells, 24 hours after treatment, stained with anti-calnexin (CNX) and anti-BiP antibodies. **C)** Expression levels of BiP, sXBP1, and CHOP genes, 48 hours after treatment, measured by RT-qPCR (one-way ANOVA)

### Treatment with the combination RBA induces oxidative stress, which is the main driver of its cytotoxic effects

ATO causes oxidative stress and ER and oxidative stress induce one another, although the underlying mechanisms are not completely clear yet^27,29^. Hence, we measured the levels of reactive oxygen species (ROS) in MOLM-13 cells treated with the combination RBA and found them increased (Figure 3A). In the presence of oxidative stress, cells trigger the oxidative stress response, mainly orchestrated by the transcription factor Nrf-2. Nrf-2 activates the transcription of genes containing an antioxidant response element (ARE) in their promoter, among which HMOX^30^. The combination RBA induces a strong oxidative stress response as demonstrated by increased expression of Nrf-2 and HMOX (Figure 3B, 3C, 3D and Supplemental Figure 2). It is important to note that ATO and all its combinations augment HMOX expression, but the combination RBA induces its highest levels. Furthermore, after 48 hours, cells treated with ATO and Btz in double combination are recovering from oxidative stress whereas those treated with RBA are not. Accordingly, the combination RBA causes the most evident alterations in mitochondria, as shown by disrupted cristae morphology observed by TEM, a sign of mitochondria suffering (Figure 3E). Indeed oxidative stress damages mitochondria by affecting the mitochondrial respiratory chain, altering membrane permeability and calcium homeostasis, and weakening the mitochondrial defense system^31^. These data indicate that ATO alone, or in double combination with RA or Btz induces oxidative stress but to a minor extent than the triple combination. Proteotoxic stress resulting from RA-mediated ER activation and the hindrance of ERAD by Btz generates oxidative stress. In RBA treatment this oxidative stress and that due to ATO converge. Very importantly, the reduction of oxidative stress by the addition of the reducing agent N-acetylcysteine is sufficient to render MOLM-13 cells resistant to all the combinations of RA, Btz, and ATO, including the triple one (Figure 3F). This is not surprising since we have already demonstrated that the reduction of oxidative stress allows AML cells to better cope with proteotoxic stress and survive^19^. Furthermore, a pivotal work by Han and colleagues shows that the main driver of cell death upon ER stress is the generation of oxidative stress^32^.

**Figure 3.**
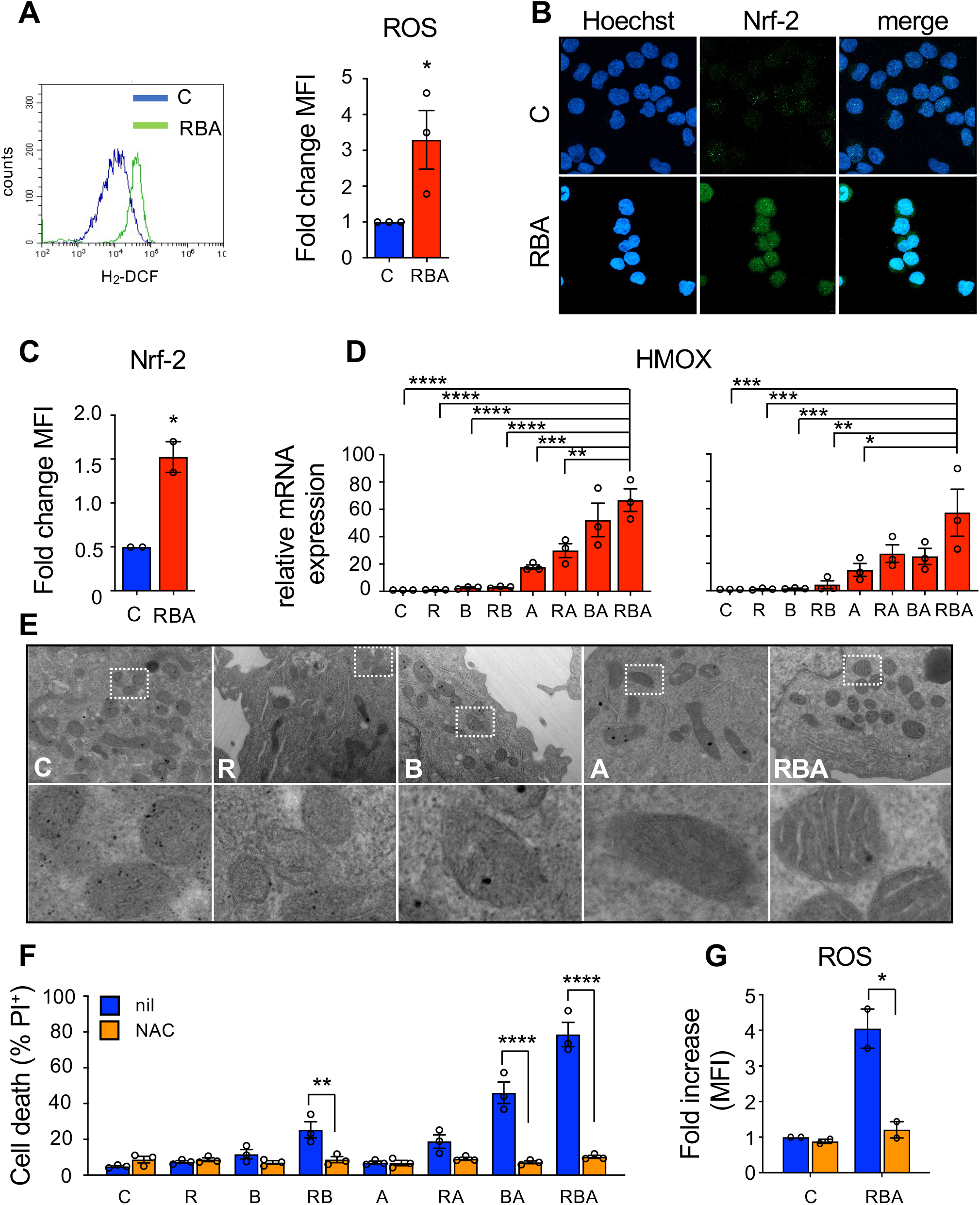
Oxidative stress is mostly responsible for the cytotoxic effect of the combination on MOLM-13 cells. **A)** ROS levels were quantified in MOLM-13 cells by flow cytometry, upon the incorporation of the oxidation-sensitive dye CMH_2_-DCFDA, 48 hours after RBA treatment (unpaired Student’s T test). **B)** Confocal microscopy images of MOLM-13 cells stained with an anti-NRF-2 antibody. **C)** NRF-2 expression in MOLM-13 cells, 24hrs after treatment, measured by flow cytometry (unpaired Student’s T test). **D)** HMOX1 gene expression levels 24 hours (on the left) and 48 hours (on the right) after treatment, evaluated by RT-qPCR (one-way ANOVA). **E)** TEM analysis focused on mitochondria of MOLM-13 cells, 24 hours after treatment; areas delimited in dashed squares in the upper panels are magnified in the lower panels. **F)** MOLM-13 cell death was measured by PI exclusion assay, 72 hours after treatment with RBA, in the absence or in the presence of 20mM N-acetyl-cysteine (left, two-way ANOVA). ROS measurement, in cells treated with or without NAC, for 48 hours (right, two-way ANOVA).

### Murine bone marrow stromal cells protect MOLM-13 cells from the toxic effects of the combination RBA, but ascorbic acid overcomes this defense

The BM niche is a heterogenous microenvironment in which normal HSCs reside and communicate with other cell populations through both direct contact and paracrine signals^33^. This crosstalk is essential for determining HSC fate and maintaining the balance between self-renewal and differentiation. Likewise, malignant LSCs interact with the cells populating the niche, altering the composition of the microenvironment to deceive the immune system, proliferate indefinitely, and invade the BM and other organs. BMSC play a main role in supporting AML progression and resistance to therapies by a variety of mechanisms^16^, most still to be elucidated, among which enhancing AML cells’ antioxidant defenses^17^. Thus, it was essential to assess the efficacy of the combination RBA on AML cells in their presence. To this aim, we co-cultured MOLM-13 cells with mouse primary BM mesenchymal stem cells/BMSC freshly isolated or with the mouse BMSC cell line MS-5. We chose murine cells in the perspective to use the combination in murine models of human leukemia (see below). In both cases, the stromal cells completely protected MOLM-13 cells from the effects of the combination RBA (Figure 4A). Since oxidative stress plays a pivotal role in the toxicity of the combination RBA, and based on literature we expected that BMSC would enhance AML cells’ antioxidant defenses, we looked for an adjuvant to further increase oxidative stress, without causing general toxicity. We chose to use ascorbic acid, or vitamin C. Ascorbic acid (ASC) is known as an anti-oxidant, which is true at micromolar concentrations. However, different studies show that at millimolar concentrations it causes oxidative stress and it has been used in clinical trials as an adjuvant of chemotherapy, without generating toxic effects^34,35^. Addition of ASC to the combination RBA completely rescued its cytostatic effects and partially its cytotoxic effects on MOLM-13 cells (Figure 4B upper panels) without affecting the viability or proliferation of MS-5 stromal cells (Figure 4B lower panels). ROS levels resulted low in MOLM-13 cells 72hrs after treatment with RBA or RBA plus ASC, showing just a slight increase in the cells treated with ASC alone (Figure 4C). However, the extensive increase of HMOX expression indicates the presence of high levels of oxidative stress in MOLM-13 treated with the combination RBA plus ASC (Figure 4D). A very strong antioxidant response could explain the low amounts of ROS. On the contrary, HMOX is only slightly up-regulated in MOLM-13 treated with ASC alone. Importantly, a comparison of HMOX expression levels in MOLM-13 cells treated with the combination RBA in mono-culture or in co-culture with MS-5, argues that in the latter condition they undergo lower levels of oxidative stress (compare Figure 3D left panel and Figure 4D), accordingly to the inefficacy of the treatment. Treatment with the combination RBA or with ASC alone partially affects MOLM-13 ER homeostasis, as indicated by the reduction of the expression of BiP and sXBP-1 (Figure 4D). However, their down-regulation is much stronger in cells treated with the combination RBA plus ASC. Altogether these data indicate that MS-5 stromal cells protect MOLM-13 from the combination RBA or ASC alone reducing the levels of oxidative stress and preserving the functions of the ER, so the cells are only partially affected by the treatments and just slow down proliferation (Figure 4B). In this context, it is important to note that the concentration of ASC we used is dramatically toxic for MOLM-13 cells in mono-culture (data not shown). Addition of ASC to the combination RBA repristinates amounts of oxidative stress high enough that cannot be recovered by stromal cells and that worsen proteotoxic stress in MOLM-13 cells. Consequently, these undergo cell death and arrest of proliferation. Importantly, we observed that medium conditioned by a co-culture of MS-5 with MOLM-13 is sufficient to defend MOLM-13 cells by the combination RBA in mono-culture, but cell-cell contact is necessary to protect from ASC and RBA plus ASC (Figure 4E).

**Figure 4.**
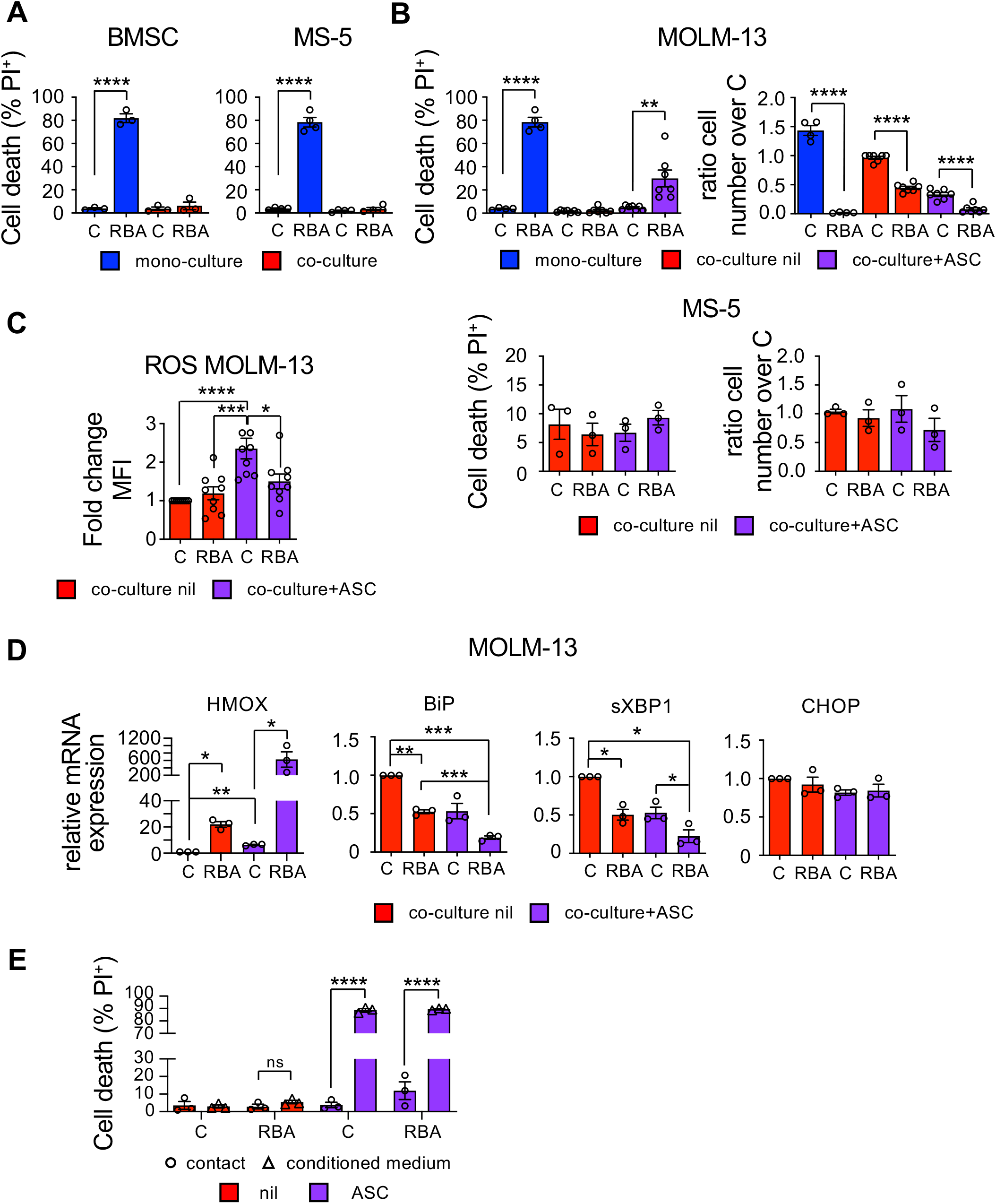
Murine bone marrow stromal cells nullify the effect of the combination RBA on MOLM-13 cells, but ascorbic acid restores it. **A)** PI exclusion assay showing cell death of MOLM-13 leukemic cells treated with 10nM RA, 2.25nM Btz and 500nm ATO (RBA) for 72 hours, in mono-culture (blue) or co-culture (red) with primary BMSC and MS-5 stromal cell line, (left and right panel respectively, one-way ANOVA). **B)** Cell death (left) and cell proliferation (right) of MOLM-13 cells (upper panels) and MS-5 cells (lower panels) treated in co-culture with RBA and 4.5mM ascorbic acid (ASC), for 72 hours (one-way ANOVA). MOLM-13 cells, treated in mono-culture, are shown as control (blue bars). **C)** MOLM-13 cells were treated in co-culture with MS-5 cells as in (B), and ROS levels were measured after 72 hours. **D)** HMOX, BiP, sXBP1, and CHOP mRNA expression levels of MOLM-13 cells, treated for 24 hours as in (B) (one-way ANOVA) **E)** MOLM-13 and MS-5 cells were treated with the combination RBA and ASC as in (B) in co-culture. After 24 hours, the conditioned medium (CM) was collected and used to treat MOLM-13 cells in mono-culture. The graph shows cell death of MOLM-13 cells treated for 48 hours with CM (one-way ANOVA).

### Mouse stromal cells treated in co-culture with MOLM-13 cells activate the oxidative stress response, undergo disruption of the actin cytoskeleton, and relocalize YAP in the cytosol

To investigate the mechanisms underlying the protection provided to AML cells, we assessed the effects of the treatments on MS-5 cells in co-culture with MOLM-13 cells. Interestingly, we observed that MS-5 cells activate their anti-oxidant defenses as indicated by up-regulation of Nrf-2, slightly upon treatment with RBA alone and more with ASC or RBA plus ASC (Figure 5A); ROS levels decrease accordingly (Figure 5B). The UPR is not perturbed by the treatments in MS-5 since we did not detect changes in BiP or sXBP-1 expression, confirming that MS-5 cells are not as affected as MOLM-13 (Figure 5C). Although, as shown above, we found no differences in the viability or proliferation of MS-5 cells (Figure 4B), we noticed modifications in their morphology (data not shown). This prompted us to analyze the structure of the actin cytoskeleton and surprisingly we found that the treatment with RBA plus ASC in co-culture caused significant disruption, with evident shortening of actin microfilaments (Figure 5D left panel and Supplemental Figure 3). As mentioned above, leukemic cells alter BMSC functions to their advantage, thus we wondered if actin cytoskeleton modifications were simply due to the treatments or because of the crosstalk with MOLM-13 in response to the treatments. Importantly, MS-5 cells cultured and treated alone do not show any change in their actin cytoskeleton, suggesting that this phenomenon is a consequence of the interaction with MOLM-13 in response to stress (Figure 5D right panel).

**Figure 5.**
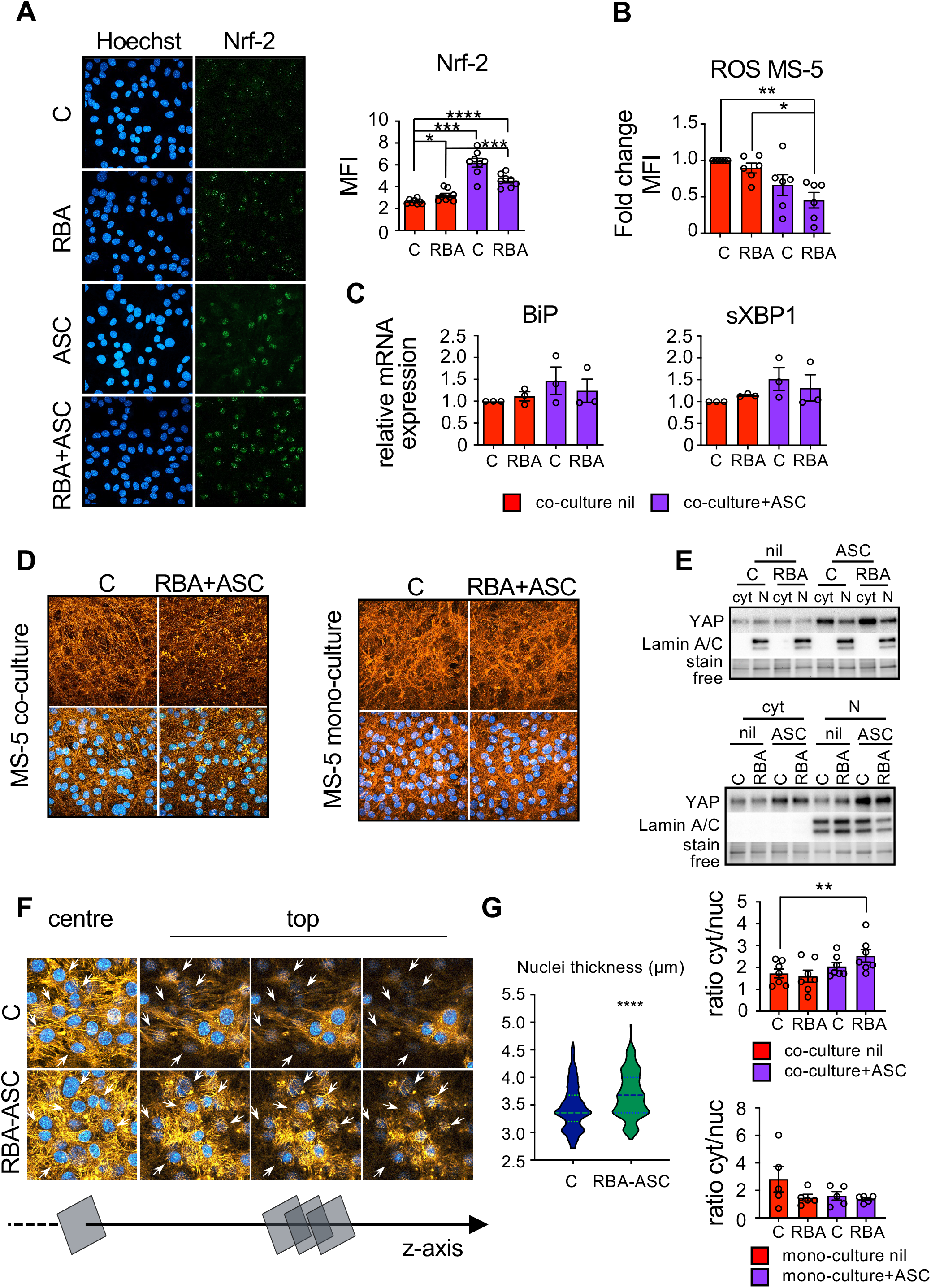
BMSC activate the oxidative stress response, undergo cytoskeletal rearrangements and re-localize YAP in the cytosol, when treated with the combination RBA plus ASC in coculture with leukemic cells. MS-5 cells were treated in co-culture with MOLM-13 for 72 hours, then analyzed. **A)** Evaluation of Nrf-2 expression by confocal microscopy. The graph on the left reports the quantification of Nrf-2 mean fluorescence intensity (n=8 fields ± S.E.M., one-way ANOVA). **B)** ROS measurement by flow cytometry (one-way ANOVA). **C)** Assessment of BiP and sXBP1 mRNA expression levels 24 hours after treatment (one-way ANOVA). **D)** Representative images of MS-5 cells stained with phalloidin, to detect F-actin (orange). Cells were counterstained with Hoechst to identify nuclei (blue). The right panels show the same analysis, but MS-5 cells were treated with the combination RBA plus ASC in the absence of the MOLM-13 cells. **E)** YAP protein levels in MS-5 cells, treated in co-culture with MOLM-13 cells (upper panel) or in monoculture (lower panel), were evaluated by western blot analysis, upon nucleus-cytosol fractionation. Lamin A/C was used as nuclear marker and stain free protein detection technology (BioRad) for total protein normalization. The graphs report the average ratio of YAP amount in the cytosol over that in the nucleus (paired Student’s T test). **F)** Single confocal images of a Z stack, taken in the center or in the apical portion of MS-5 cells, stained as in (D), to examine the actin cap. **G)** The violin plot reports the measurements of nuclear thickness, obtained by the confocal analysis shown in (F).

It is well assessed that transformations in the mechanical environment and actin cytoskeleton regulate the transcriptional activator YAP through Hippo pathway dependent and independent mechanisms^36^. In particular, when the actin cytoskeleton is well developed, with an abundance of actin microfilaments and stress fibers, YAP is localized in the nucleus and is transcriptionally active. On the contrary, when the actin cytoskeleton is disrupted (for example when cells grow in soft extracellular matrix (ECM) or onto small surfaces or at high density) YAP is relocalized in the cytosol and its target genes are silenced^37,38^. Thus, we measured the ratio between cytosolic and nuclear YAP in MS-5 cells upon treatment in co-culture with MOLM-13. We found that YAP is partially relocalized in the cytosol of cells treated with RBA plus ASC, indicating downregulation of its activity (Figure 5E upper panel). Accordingly to our observations relative to the actin cytoskeleton, MS-5 cells treated alone do not relocalize YAP in the cytosol, meaning that cytoskeletal disruption and YAP modulation are due to the interaction with MOLM-13 cells (Figure 5E lower panel). The regulation of YAP localization and activity in relation to the actin cytoskeleton is quite complex, regulated at multiple levels, and not completely elucidated yet^36,39,40^. Recent works show that YAP transcriptional activity and actin structure could be involved in a feedback loop regulating the integrity of the nuclear envelope^41^. Moreover, it has been shown that a flatter shape of the nucleus, which is maintained by a well-developed actin cap^42^, favors YAP entry into the nucleus^43,44^. Thus, we analyzed by confocal microscopy the structure of the nuclear actin cap of MS-5 cells treated with RBA plus ASC in the presence of MOLM-13 cells and found that it is disrupted, characterized by short and disordered actin filaments, contrary to untreated cells in which the actin filaments on top of the nuclei are elongated and orderly integrated into the actin network (Figure 5F). As a consequence, the nuclei thickness of treated cells is significantly augmented. The graph shows the expansion of subpopulations of MS-5 cells presenting nuclei with heightened thickness and a decrease in those with shorter one (Figure 5F and Supplemental Figure 4).

Altogether our data suggest that the crosstalk between AML cells and stromal cells, in the presence of stress generated by the combination RBA plus ASC, induces changes in the stromal cells that activate the oxidative stress response and undergo disruption of the actin cytoskeleton. As a consequence, part of the stromal cells presents rounder nuclei which could be one of the mechanisms underlying partial YAP inactivation by cytosolic relocalization.

### The combination RBA plus ascorbic acid prolongs the life span of a murine model of human AML

To assess the efficacy, and possible toxicity, of the combination RBA plus ASC in a murine orthotopic model of human AML *in vivo,* we exploited the NSG mouse model engrafted with MOLM-13 cells. We observed a significant increase in the life span of mice treated with RBA plus ASC (Figure 6A) compared to engrafted, untreated animals, without differences in behavior, body weight, and tissue morphology of spleen, liver and kidney (Supplemental Figure 5A). Importantly, analysis of the femurs’ bone marrow at the time of sacrifice evidenced significantly lower leukemia engraftment in treated mice (Figure 6B and Supplemental Figure 5C). Furthermore, most of the MOLM-13 cells recovered from the treated mice show extensive vacuolation (but the murine granulocytes do not), contrary to the cells recovered from the control mice, in which just a small minority of MOLM-13 cells presents such vacuoles. These findings indicate that the treatment RBA plus ascorbic acid specifically hits AML cells in the BM *in vivo*, without evident general toxicity.

**Figure 6.**
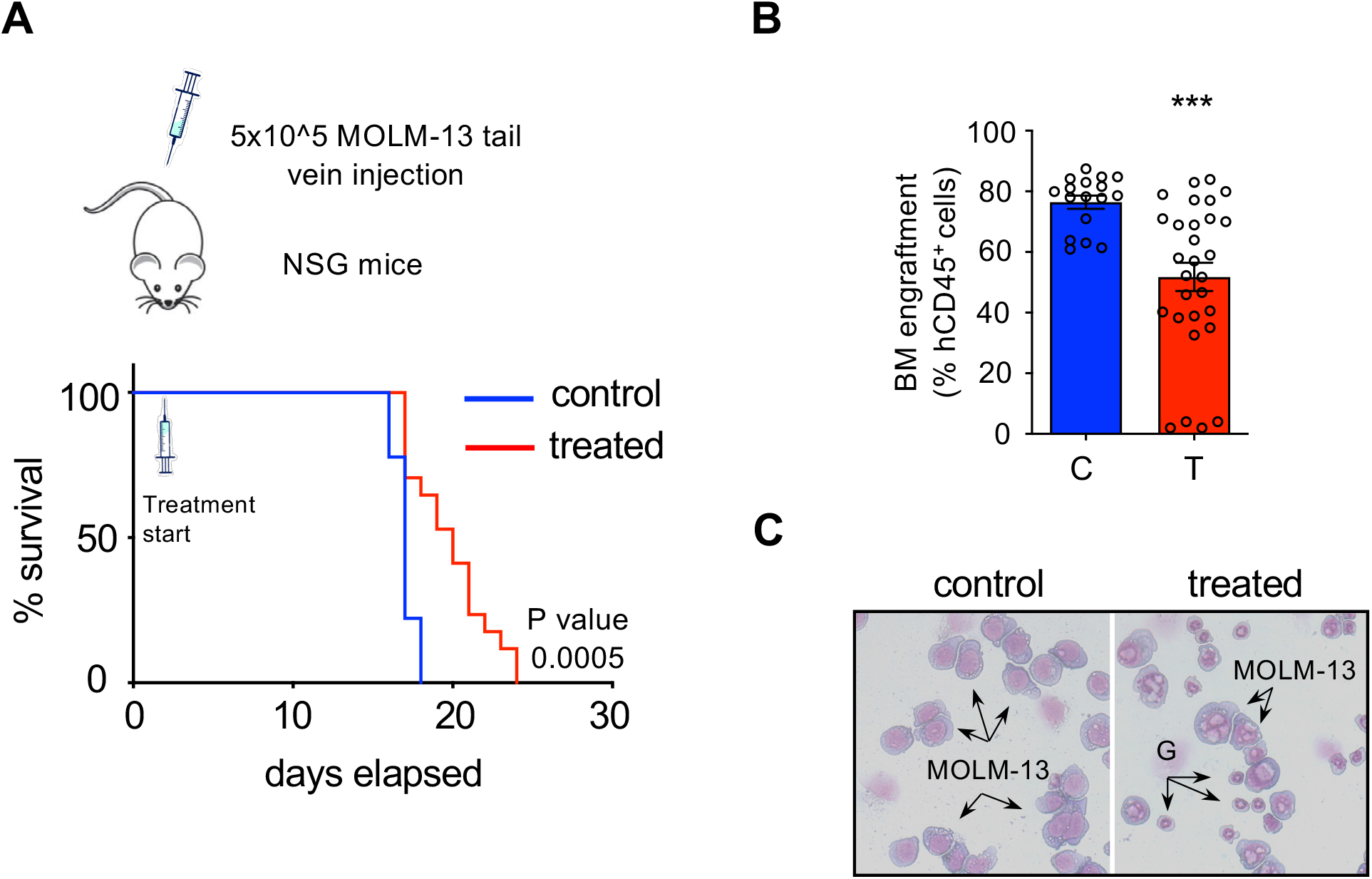
The combination of RBA plus ASC acid significatively prolongs life-span of NSG mice engrafted with human FLT3-ITD^+^ MOLM-13 cells. **A)** Kaplan-Meyer survival plot showing OS of NSG mice engrafted with MOLM13 leukemic cell line and treated with vehicle or RBA+ASC, starting at day 2 after cell injection. (control=9 and treated=17 from two independent experiments) **B)** The treatment with RBA plus ASC reduces the percentage of leukemic cells in the BM of engrafted mice. Both control and treated mice were sacrificed when showing posterior limb paralysis. Bone marrow was flushed by femurs and purified mononucleated cells were analyzed by flow cytometry after staining with an anti-human CD45 antibody (unpaired Student’s T test) **C)** Wright-Giemsa staining of the cells collected from mice bone marrow shows extensive vacuolation of MOLM-13 cells isolated from the BM of treated mice present to a much lesser extent in those obtained from control ones. G indicates murine granulocytes that show no vacuolation.

## Discussion

The frontline of AML cure is focusing on molecular targeted therapy although, at present, the development of resistance and consequent relapses remain major issues^45^. Clonal selection is one of the main obstacles to a successful cure^15^. The most promising strategy to restrict this phenomenon is to target different oncogenic/survival pathways in combination. Protein homeostasis is essential for cell survival and leukemia cells depend on adaptive stress responses^18,46,47^. We have already demonstrated that FLT3-ITD^+^ AML cells are particularly sensitive to proteostasis unbalance, also because this mutated protein is unfolded and retained in the ER by its quality control systems^19^. To increase the translational potential of our strategy we use here the proteasome inhibitor bortezomib to induce proteotoxic stress, in combination with retinoic acid and arsenic trioxide, since, as the last two, Btz is already available in clinical practice. The use of a combination of drugs at low doses targeting different pathways, all inducing proteotoxic stress, should increase specificity and decrease off-target toxicity. Indeed, both *in vitro* and *in vivo* we observe strong effects on FLT3-ITD^+^ AML cells but not on other cell types or organs.

In addition to clonal selection, the complex framework of the disease comprises crosstalk between AML cells and the BM niche upon therapy administration^16^. In fact, we observed that BMSC protect AML cells from the cytotoxic effects of the combined treatment. Our findings, indicating that oxidative stress is at the basis of the efficacy of the combination RBA, suggested using ascorbic acid as an adjuvant to exacerbate it and overcome resistance. In this context, it is clear the importance of understanding the mechanisms underlying AML-stromal cell crosstalk. AML cells and BMSC exchange cytosolic content by tunneling nanotubes (TNT), which are formed by AML cells and take contact with BMSC^48^. These are sustained by an actin structure^49^, mediate the transfer of mitochondria from BMSC to AML cells and increase the latter anti-oxidant defenses and resistance to chemotherapy^17,48^. Our study reveals that the combination RBA plus ascorbic acid disrupts the structure of the BMSC actin cytoskeleton. Although the molecular mechanism must still be investigated, this could play a main role in the ability of this treatment to overcome the resistance of AML cells provided by the BMSC. It is important to underline that actin cytoskeleton disruption in MS-5 is not merely the effect of the stress generated by the combination RBA plus ASC, but it is a consequence of the interaction between MOLM-13 and MS-5 cells in condition of stress. Notably, actin cytoskeleton structure is closely linked to the activity of the transcriptional coregulator YAP, which plays a main role in development, tissue homeostasis, and cancer. We observe that, upon treatment of the co-cultures, MS-5 cells partially relocalize YAP in the cytosol, thus restricting its activity. To our knowledge, this is the first time in which YAP is found to be involved in the crosstalk between AML and BM stromal cells.

Many critical questions remain to be answered and are already matters of investigation in our lab. The first concerns the meaning of YAP relocalization in MS-5 upon treatment with RBA plus ASC. It will be of paramount importance to understand the role of YAP in the leukemic niche since manipulation of its activity in the BM environment could become an important therapeutic strategy. The second relevant issue to be investigated regards the mechanisms determining actin cytoskeleton disruption. It has been shown that AML cells promote mitochondrial transfer from BMSC, via TNT, by increasing their levels of oxidative stress via NOX2 activity^48^. The combination RBA plus ASC dramatically alters the redox balance of AML cells and this could have consequences on actin remodeling in BMSC, which is involved in TNT formation. Another possibility could be that the interaction of stromal cells with leukemic cells undergoing stress causes ECM rearrangements, rendering it “softer”, which would induce actin cytoskeleton disruption and YAP cytosolic relocalization in stromal cells^36,40^. The BM ECM is modified in AML and myelodysplastic syndrome^50,51^, and AML cells produce a different secretome than HSC or normal progenitors^51,52^. The combination RBA plus ASC could interfere with this equilibrium that provides advantages to AML cells, with consequences on stromal cell mechanotransduction.

On the whole, we report the possibility of specifically targeting FLT3-ITD^+^ AML cells by inducing proteotoxic stress with the combination RBA plus ascorbic acid, all drugs already in use in clinical practice in different settings. Furthermore, we show that the inability of BMSC to protect AML cells from the combination correlates with actin cytoskeleton disruption and YAP relocalization into the cytosol, suggesting for the first time that this mechanism could be involved in the protective functions of the BMSC and opening to the identification of novel therapeutic targets.

## Acknowledgments

The research leading to these results has received funding from: AIRC IG 2018-ID. 21406 project; Fondazione GIMEMA “Fondo per le Idee” 2018; Istituto Pasteur Italia-Fondazione Cenci Bolognetti; ‘Progetti Ateneo’ Sapienza University of Rome; PRIN 2017-Prot. 2017TATYMP_003; NextGenerationEU” DD. 3175/2021 E DD. 3138/2021 CN_3: National Center for Gene Therapy and Drugs based on RNA Technology Codice Progetto CN 00000041.

## Authorship Contributions

FL, MS and SM performed and analyzed the experiments and contributed to the experimental design. CT and AI contributed to the experiments. TO, MD, and ST characterized and provided leukemic bone marrow samples and contributed to the characterization of leukemic cells recovered from engrafted mice. MM and RC collaborated with MS for the *in vivo* experiments. SM, GF, SAN and VP provided the expertise and the technology for the TEM analysis. LT and MTV contributed to the experimental design and critical reading of the manuscript. SM and FF planned the research and wrote the paper. All authors read and approved the final manuscript.

## Disclosure of conflicts of interest

The authors declare no competing interests.

## Notes

### Competing Interest Statement

The authors have declared no competing interest.

