## Supplemental material for "Retinoic acid and proteotoxic stress induce AML cell death overcoming stromal cell protection"

**Supplemental methods**

**Cell lines and primary cell culture and treatment.**

Human acute myeloid leukemia cell lines MOLM-13, MV4-11, MOLM-14, ML-2, OCI-AML2, OCI-AML3, HL60 and NB4 were obtained by DSMZ (Leibniz Institute, Germany) and cultured in RPMI 1640 medium supplemented with 1% penicillin/streptomycin and 10% heat-inactivated FBS (Gibco, ThermoFisher Scientific, Waltham, MA, USA). AML bone marrow samples were collected at diagnosis in the Department of Biomedicine and Prevention at the University of Rome Tor Vergata after obtaining written informed consent to the study from all patients. The study was approved by the IRB of Policlinico Tor Vergata Rome. The murine mesenchymal stem cell line MS-5 was obtained by DSMZ and cultured in MEM Alpha medium with ribonucleosides and deoxyribonucleosides (Gibco, ThermoFisher Scientific), supplemented with 1% penicillin/streptomycin, 20% heat-inactivated FBS and 2-mercaptoethanol 100uM (Sigma-Aldrich, St. Louis, MO). Primary murine bone marrow stromal cells were obtained from three healthy mice. After sacrifice, the femurs were flushed with a syringe in order to discard hematopoietic cells. The bones were cleaned and crushed in a mortar, then the bone matrix was digested by collagenase for 30-40 minutes. Isolated cells were seeded in a medium with 20% FBS enriched with growth factors: 100uM 2-ME, 1X NEAA, N2, B27, 10ng/ml EGF, 40ng/ml IGF, FGF, 40ng/ml PDGF, Oncostatin M (Peprotech, Rocky Hill, NJ, USA). After the first passage, only 2-ME, NEAA, IGF, and EGF were maintained.

All the cultures were maintained at 37°C in a humified atmosphere containing 5% $CO_2$.

MOLM13 cells were treated with 10nM Retinoic Acid (R), 2.25nM Bortezomib (B) and 500nM Arsenic Trioxide (A) bought from Sigma-Aldrich, alone or in combination, as described in the text. In order to attenuate oxidative stress, cells were pre-incubated for 24 hours with N-Acetyl cysteine (NAC) 20mM (Sigma-Aldrich) at pH 7.4 and then treated with the above-mentioned drugs, adding again also NAC 20mM. Primary cells were thawed and the leukemic stem cells ($CD34^+$) were isolated by positive selection with CD34 MicroBead Kit UltraPure human (Miltenyi Biotec, Bergisch Gladbach, Germany), following the manufacturer's instructions. Both $CD34^+$ and $CD34^-$ cells obtained from the separation were seeded at a concentration of 10.000 cells/ml in StemSpan Leukemic Cell Culture Kit (STEMCELL Technologies, Vancouver, Canada). After waiting 7 days for cell amplification, they were treated with 10nM RA, 3nM Btz, and 1mM ATO, then incubated at 37°C with 5% $pCO_2$. For the co-culture experiments, the MEM Alpha medium in dishes of murine stromal cells almost confluent was changed with RPMI, and 24 hours later this was removed in order to add fresh RPMI containing

MOLM13 leukemic cells and drugs. For the co-culture experiments, Ascorbic acid (by Sigma-Aldrich) was used at 4.5mM final concentration.

**Cell death, cell cycle, morphologies, and TEM.**

Leukemic cell lines seeded at 70.000 cells/ml or 100.000 cells/ml depending on their doubling time, were treated with different drugs as indicated in the text. At the indicated time points cells were harvested and stained with 10ug/ml propidium iodide (Sigma-Aldrich, St. Louis, MO, USA). In the co-culture experiments, the same procedure was performed for stromal cells upon detachment with 0.05% trypsin/0.53mM EDTA (by Corning, Corning, NY, USA). Both cell death and proliferation were analyzed by flow cytometry (Cytoflex Beckman Coulter, Life Sciences, Brea, CA, USA) and CytExpert v2.2 software, (Beckman Coulter). 24 hours and 48 hours after treatments, 0.5 $x10^6$ cells were washed with PBS and then fixed O/N with 70% ethanol at +4°C. Then, cells were incubated for at least 3 hours with a propidium iodide solution with RNase. After this step, cells were analyzed by flow cytometry for their DNA content.
In order to analyze cell morphology, 72h hours after treatment about 300.000 leukemic cells were spotted on a slide by using a Shandon Cytospin 3 Centrifuge (ThermoFisher Scientific, Waltham, MA, USA). Cells were stained with Giemsa stain, modified solution, following the manufacturer's instruction (Sigma-Aldrich). Pictures of an appropriate number of representative fields from each sample were acquired by using Zeiss Axiocam 503 color. Zen blue software (Zeiss) and Adobe Photoshop (Adobe Systems) were used for the processing of images.
In order to analyze the intracellular structures upon treatments, about 30-40x10^6 cells for each condition were collected 24 hours after drug administration and centrifuged at 300xg for 5 minutes. The cell pellet was fixed with 2.5% glutaraldehyde (SIC, Rome, Italy) in 0.1 M PBS for 2 days at 4 °C and then rinsed with PBS. Then the samples were post-fixed using 2% osmium tetroxide (Agar Scientific, Stansted, UK) for 2 h. The specimens were dehydrated by exchange with ethanol (30%, 70%, 95%, 100% v/v x3), immersed in propylene oxide (BDH Italia, Milan, Italy) for solvent substitution and embedded in epoxy resin Embed-812 (SIC, Rome, Italy). Ultrathin sections (70-80 nm) were obtained using a Leica EM UC6 ultramicrotome and collected on 100 - mesh copper grids (Assing, Rome, Italy) and stained with a mix of lanthanides solution (Uranyless, Electron Microscopy Sciences) and lead citrate. Imaging was performed using a Zeiss EM10 trasmission electron

microscope set at an accelerating voltage of 60 kV, and images were captured using an AMT CCD digital camera (DEBEN UK Ltd).

**RNA extraction and RT-qPCR.**

Total RNA was isolated by TRIzol reagent (Invitrogen, ThermoFisher Scientific, Waltham, MA, USA) following the manufacturer's instructions. After quantification and quality check by measuring OD at 230, 260, and 280nm, 250ng of total RNA were reverse-transcribed with the High-Capacity RNA to cDNA kit (Applied Biosystems, ThermoFisher Scientific, Waltham, MA, USA) in a final volume of 10μl, then diluted 1:5. 1μl of the dilution was used for each amplification. Intercalant dye-based Real-Time PCR was performed by using PowerUp SYBR Green Master Mix (Applied Biosystems, ThermoFisher Scientific, Waltham, MA, USA) in a final volume of 10μl. The reactions, all performed in duplicates, were carried out by using custom oligonucleotides, designed as follows:
H3 (FW: GTGAAGAAACCTCATCGTTACAGGCCTGGT - RW: CTGCAAAGCACCAATAGCTGCACTCTGGAA);
hBiP (FW: TAGCGTATGGTGCTGCTGTC - RW: TTTGTCAGG GGTCTTTCACC);
hCHOP (FW: TGGAAGCCTGGTATGAGGAC - RW: TGTGACCTCTGCTGGTTCTG);
spliced XBP1 (FW: GAGTCCGCAGCAGGTGC - RW: TCCTTCTGGGTAGACCTCTGGGAG). H3 was used as the reference gene. HMOX1 expression levels were evaluated by the PrimeTime Std qPCR Assay (Integrated DNA Technologies, Skokie, Illinois, USA) containing the following primers
HMOX1 (FW: TCATGAGGAACTTTCAGAAGGG – RW: TGCGCTCAATCTCCTCCT) and probe (/56-FAM/TGGCCTCCC/ZEN/TGTACCACATCTATGT/3IABkFQ/).
For these reactions, the PrimeTime Std qPCR Assay for GAPDH was used as reference:
GAPDH (FW: ACATCGCTCAGACACCATG – RW: TGTAGTTGAGGTCAATGAAGGG) and probe (/56-FAM/AAGGTCGGA/ZEN/GTCAACGGATTTGGTC/3IABkFQ). The reactions were performed with the TaqMan Universal PCR Master Mix reagent (Applied Biosystems, ThermoFisher Scientific, Waltham, MA, USA). All qPCRs were performed by the QuantStudio 7 Flex Real-Time PCR (Applied Biosystems, ThermoFisher Scientific, Waltham, MA, USA) and analyzed with QuantStudio Real-Time PCR Software v1.3 (Applied Biosystems, ThermoFisher Scientific, Waltham, MA, USA).

**Immunofluorescence and flow cytometry analysis.**

About 200-400.000 MOLM-13 cells were harvested at the time points indicated in the figures, washed, and fixed in 4% PFA for 10 minutes. Fixed cells were washed and permeabilized with 0.1% TritonX100 in PBS-BSA 1% for 10 minutes. Cells were stained with primary antibodies in PBS-BSA 1% for 30 minutes at room temperature. The primary antibodies used are: Calnexin (C5C9) Rabbit mAb (#2679, Cell Signaling Technology, Danvers, MA, USA), 1:100; BiP/GRP78 Mouse mAb (610978, BD Transduction Laboratories, Franklin Lakes, NJ, USA), 1:100; NRF2 (D1Z9C) XP Rabbit mAb (#12721, Cell Signaling), 1:500. After washes in PBS-BSA 1%, cells were incubated with Alexa Fluor conjugated secondary antibodies: Goat anti-Rabbit IgG (H+L) Highly Cross-Adsorbed Secondary Antibody, Alexa Fluor 488 (ThermoFisher Scientific), 1:500; Goat anti-Mouse IgG (H+L) Cross-Adsorbed Secondary Antibody, Alexa Fluor 555 (ThermoFisher Scientific), 1:500. F-actin was stained with Rhodamine-Phalloidin R415 (ThermoFisher Scientific) 1:50 and human CD45 was detected by a FITC-CD45 antibody (Biolegend, San Diego, CA, USA). After 30 minutes cells were washed and incubated with 1:1000 Hoechst 33342 solution. MOLM-13 cells were washed and resuspended in 15μl of PBS, then 4μl were spotted on a slide and let dry. The same procedure was used for MS-5 cells, but these were seeded on glass slides at the beginning of the experiment. In both cases, the slides were mounted with Vectashield mounting medium (Vector Laboratories, Burlingame, CA, USA) and closed with a cover glass. Confocal images were acquired by Zeiss LSM 900 with Airyscan 2, equipped with ZEN 3.2 Blue Edition software. Stromal cell nuclei thickness on z-axis was assessed by acquiring sections with 0.16uM step and measuring the 3D distance in the Orthogonal section, using ZEN 3.2 Blue Edition software (Supplemental Figure 4). Adobe Photoshop was used for the processing of images. ImageJ software was used to quantify the mean fluorescence intensity of an appropriate number of fields from each experimental condition, as indicated in the figures.

Spleen sections were stained with a FITC-CD45 antibody (Biolegend, San Diego, CA, USA) TO-PRO-3 and 1:3000 (ThermoFisher Scientific) to reveal nuclei.

For the detection of Nrf2 fluorescence by flow cytometry, about 400.000 cells were collected in round bottom 2ml tubes and fixed, permeabilized, and stained with primary and secondary antibodies as stated above. After washes of the secondary antibody, the cells were incubated with Sytox blue fluorescent dye (ThermoFisher Scientific) to exclude dead cells. Data were acquired by a Cytoflex Beckman Coulter flow cytometer, from Life Sciences. The results were analyzed with CytExpert v2.2 by Beckman Coulter software.

For ROS measurement by flow cytometry around 400.000 cells were harvested at different time points, abundantly washed and incubated with 2μM CM-$H_2$DCFDA (ThermoFisher Scientific) in pre-warmed PBS. CM-$H_2$DCFDA is a general oxidative stress indicator and it derives from $H_2$DCFDA. This molecule passively diffuses into cells, where it is cleaved by esterases. The thiol-reactive chloromethyl group is oxidized by cellular ROS, thus becoming a fluorescent adduct, and the more ROS are present the more fluorescence detected will be intense. After incubation with CM-$H_2$DCFDA at 37°C for 30 minutes in the dark, cells were washed and stained with Sytox blue fluorescent dye in order to exclude dead cells. Also in this case, samples were analyzed with Cytoflex Beckman Coulter and the results were analyzed with CytExpert software.

**Western blot.**

MS5 stromal cells obtained from the co-culture and mono-culture experiments were washed with cold PBS and lysed in the dishes in RIPA buffer containing 150mM NaCl, 10mM Hepes, 0.25% Sodium Deoxycholate, 1% NP40, and 0.1% SDS. Lysates were spun for 5 minutes at 300xg. The cytosolic fraction was collected and the residual nuclear fraction was lysed in a buffer containing 25mM Tris pH 7.5, 100mM NaCl, 3mM EDTA, 7% glycerol, and 2% SDS. After the sonication of the nuclear fraction, the protein samples were separated by SDS-PAGE, on stain free BioRad gels (BioRad, Hercules, CA, USA) and transferred to a nitrocellulose membrane. The membrane was incubated overnight with the following primary antibodies: anti-YAP Rabbit-mAb (#4912S Cell Signaling Technology) and anti-lamin (#4912S Cell Signaling Technology). Detection of the western blots was performed with ECL western blotting system (GE Healthcare, Sigma-Aldrich St. Louis, MO, USA) and images of the blots were obtained with the ChemiDoc XRS+, using the Image Lab software (Bio-Rad).

***In vivo* experiments.**

*NSG (Jax ™ NOD.Cg-Prkdcscid Il2rgtm1Wjl/SzJ)* mice (Jackson Lab, Bar Harbor, ME, USA) were injected in the tail vein with 500.000 MOLM-13 cells. Treatments started 2 days after engraftment. Retinoic acid was administered as 5 mg pellets, with a release rate of 21 days, (Innovative Research of America, Sarasota, FL, USA), inserted under the skin of the animal with a trochar. Bortezomib was administered at 0,5 mg/kg in 100µl PBS, via IP injection every 4 days for 3 weeks (5 doses in total), arsenic trioxide at 3 mg/kg in 100 µl PBS via IP injection every day for 3 weeks, and ascorbic acid at 1g/kg via IV injection. Mice were monitored for weight (every week) and behavior (every day) and were sacrificed when leukemia symptoms led to hind limbs paralysis. Spleens, livers, and kidneys were harvested and preserved at -80°C in OCT (Leica Biosystems, Wetzlar, Germany) for further histological analysis. Paraffin embedding and H&E staining were performed as standard (Cardiff, Miller and Munn. Cold Spring Harb Protoc. 2014 Jun 2;2014(6):655-8. doi: 10.1101/pdb.prot073411). Sections for spleen IF were obtained by OCT frozen tissue with a cryostat and then processed as described above. Bone marrow was flushed from the femurs, red cells lysed, and nucleated cells were diluted in 3ml PBS and passed through 70μm nylon cell strainers (Corning, Corning, NY, USA) to remove small pieces of bones and big debris. Cells were counted and 300,000 cells were processed to observe cell morphology as described above for MOLM-13. 200,000 cells were stained 30' at 4°C with FITC-hCD45 antibody 1:200 (Biolegend, San Diego, CA, USA), washed, and resuspended in 400μl PBS with 1:100 Sytox blue fluorescent dye to exclude dead cells. Samples were analyzed with a Cytoflex Beckman Coulter flow cytometer and the results were analyzed with CytExpert software.

**6.7 Statistical analysis**

All the graphs report each replicate sample with the symbol "○". The replicates are intended as biological replicates obtained in independent experiments. The error bars represent the standard error of the mean (SEM). Statistical analysis was performed by Student's T-test or ANOVA and multiple comparison tests as indicated in the legends. *P <0.05, **P <0.005, ***P <0.001, ****P <0.0001. All statistical analyses were performed by the GraphPad-Prism 8 software (GraphPad Software, La Jolla, CA, USA).

| Cell line | Genotype | Sensitive to RBA |
|---|---|---|
| **MOLM13** | t(9;11)/MLL-AF9; **FLT3-ITD** | ++++ |
| **MV4-11** | t(4;11)/MLL-AF4; **FLT3-ITD** | +++ |
| **MOLM14** | t(9;11)/MLL-AF9; **FLT3-ITD** | ++ |
| **ML2** | t(6;11)/MLL-AF6; KRAS$^{A146T}$ | + |
| **OCI-AML2** | t(6;11)/MLL-AF6 | - |
| **OCI-AML3** | DNMT3A$^{R882C}$; NRAS$^{G61}$; NPM1$^{W288Cfs*12}$ | - |
| **HL60** | del TP53; CDKN2A$^{R80Ter}$; NRAS$^{G61L}$ | + |
| **NB4** | t(15;17)/PML-RARα; KRAS$^{A18D}$; TP53$^{R248G}$ | + |

**Supplemental Table 1. FLT3-ITD$^{+}$ cell lines are the most sensitive to the combination RBA.** The cell lines listed in the table were treated with different concentrations of Btz for 72hrs to produce a dose-response curve. The concentrations causing a percentage of cell death below the threshold of 15%, which varied slightly for each cell line, was chosen to be tested in combination with 10nM RA and 500nM ATO. Cell death and cell proliferation were evaluated by flow cytometry after staining with propidium iodide.

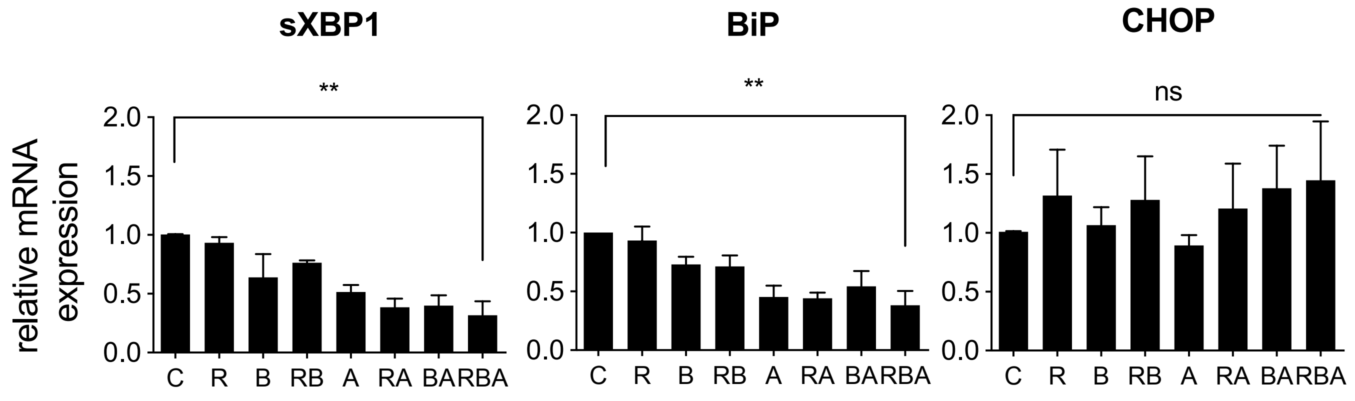


**Supplemental Figure 1. UPR gene expression upon 24hrs of treatment.** MOLM-13 cells were treated for 24hrs with the combination RBA as described in figure 1 and the expression of the UPR genes sXBP1, BiP and CHOP was measured by RT-qPCR.

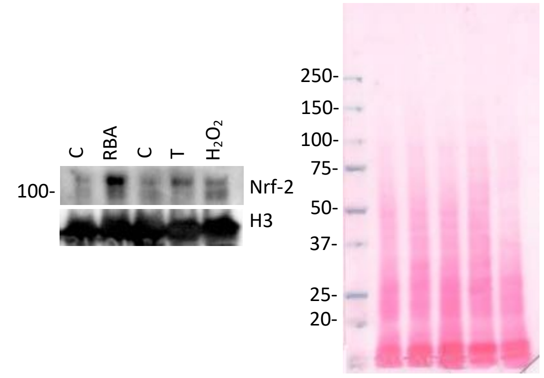


**Supplemental Figure 2. The treatment RBA induces Nrf2 protein expression and nuclear accumulation.** MOLM-13 cells were treated with vehicle (C) or 10nM RA, 2.25nM Btz, 500nM ATO (RBA) for 48hrs; vehicle (C), Tunicamycin 350nM (T) for 24hrs; $H_2O_2$ 100µM for 30'. Nuclei were isolated and lysed and proteins were analyzed by WB decorating the membrane with an anti-Nrf2 antibody followed by an anti-Histone 3 antibody. Accordingly to literature, and to our previous data[19], ER proteostasis unbalance, induced by Tm, causes oxidative stress and Nrf-2 nuclear accumulation. $H_2O_2$ was used as a control of oxidative stress due to increase in cellular ROS.

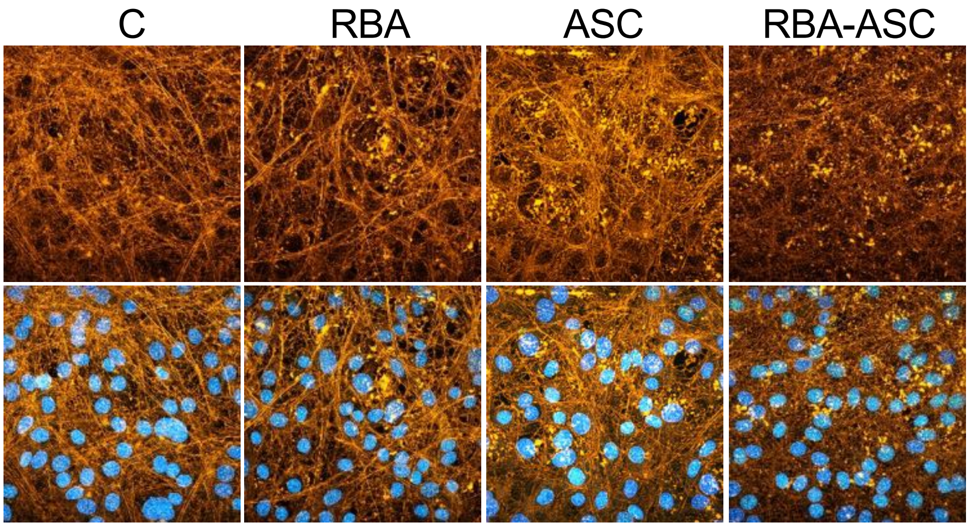


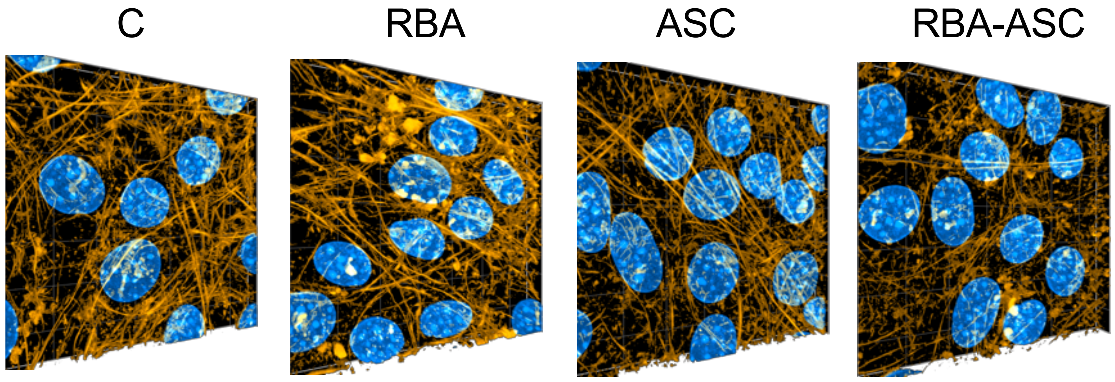

**Supplemental Figure 3. The treatment RBA plus ASC causes actin cytoskeleton disruption.** MS-5 cells were treated with the combination RBA plus ASC, in co-culture with MOLM-13, for 72 hours,. MOLM-13 cells were removed, and MS-5 cells were stained with Alexa 555-phalloidin (orange) and Hoechst (blue) to reveal F-actin and nuclei. Images were acquired by a confocal microscope. The upper panels show a representative 2D overlay of z-stack images. Only the samples treated with RBA plus ASC exhibit evident disruption of actin filaments. The lower panels show a 3D reconstruction of the Z-stack images (about 40 focal planes) obtained by super-resolution confocal microscopy

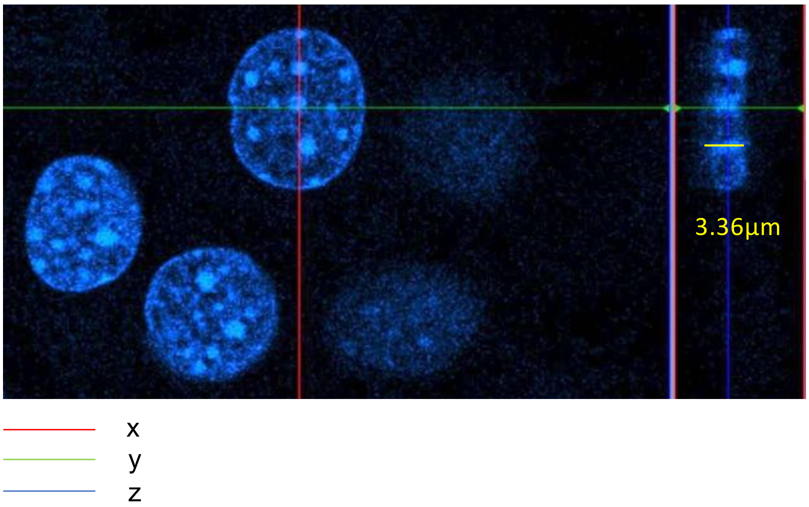


**Supplemental Figure 4. Measurement of nucleus thickness.** The same z-stack images used in Figure 3 were analyzed by the confocal microscope Zen Blue Software (Zeiss). The image reports an example of how the nuclei can be measured on the three dimensions. In Figure 6G we report measurements of thickness on the z axes, evaluated as shown here.

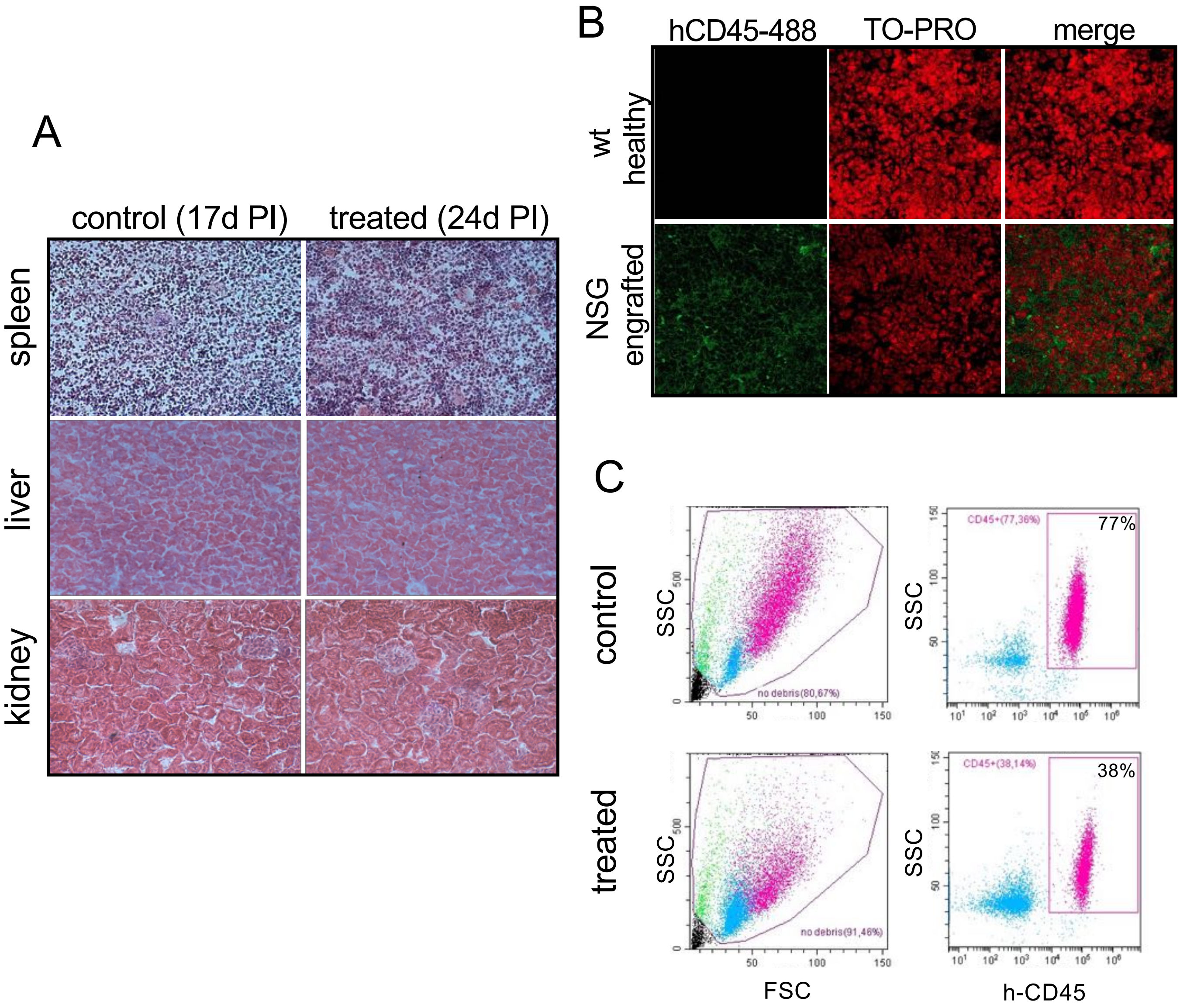


**Supplemental Figure 5. Analysis of NSG leukemic mice. A** Histological analysis of spleen, liver and kidney of NSG engrafted mice, treated or not with the combination RBA plus ASC, show no signs of toxicity due to the treatment. As described in the text all the animals were sacrificed when showing hind limbs paralysis. **B** Confocal microscopy of spleen specimen from healthy wt or MOLM-13 engrafted NSG mice demonstrates that this tissue is heavily invaded by MOLM-13 cells. Spleen specimens were harvested, snap-frozen, cut at cryostat, and stained with an Alexa 488-anti-humanCD45 to reveal MOLM-13 cells; nuclei were counterstained with the TO-PRO3 dye. **C** The panel reports an example of the flow cytometry analysis shown in figure 6C. MOLM-13 cells were detected by an Alexa 488-anti-humanCD45
